# The Longitudinal Analysis of Convergent Antibody VDJ Regions in SARS-CoV-2 Positive Patients Using RNA-seq

**DOI:** 10.1101/2022.11.02.514944

**Authors:** Kate J. Liu, Monika A. Zelazowska, Kevin M. McBride

## Abstract

The severe acute respiratory syndrome-related coronavirus-2 (SARS-CoV-2) has infected over 600 million individuals and caused over 6.5 million deaths. To understand the immune response individuals have from the SARS-CoV-2 infection, we studied the immunoglobulins against the virus’s antigens. The diversified complementarity determining region 3 (CDR3) can be used to characterize an antibody. We downloaded four public RNA-seq data sets that were collected between March 2020 and March 2022 from the Gene Expression Omnibus (GEO) in our longitudinal analysis. In total, there were 269 SARS-CoV-2 positive patients and 26 negative patients who served as a control group. Samples were grouped based on their SARS-CoV-2 variant type and/or the time they were collected. Among 629,137 immunoglobulin V(D)J sequences identified by reconstructing the V(D)J sequences, we found 1011 common V(D)Js (same V gene, J gene and CDR3 sequences in each SARS-CoV-2 positive group) shared by more than one patient in each group and no common V(D)Js were from the negative control group. In our clustering analysis, we identified 129 convergent clusters from the SARS-CoV-2 positive groups. One of these convergent clusters matched the protein sequence of crystal 3D structures of the antibodies against SARS-CoV-2 in the Protein Data Bank (PDB). In our longitudinal analysis between the Alpha and Omicron variant, we found 2.7% of common CDR3s were shared although the longitudinal profiling of common V(D)Js was variant specific. Although diverse immunoglobulin profiles were observed, the convergence of common V(D)Js suggests that there exists antibodies with similar antigenic specificities across patients in different groups over various stages of the pandemic.

## 1. Introduction

When humans are infected with severe acute respiratory syndrome-related corona-virus-2 (SARS-CoV-2), the immune system will naturally generate antibodies with binding specificity to SARS-CoV-2 and these antibodies can be detected in the blood serum of patients [1, 2]. The convergent antibodies that neutralize SARS-CoV-2 play an important role in fighting the severity of the coronavirus disease-2019 (COVID-19) [3, 4]. The characterization of antibodies will help elucidate the mechanisms of immune response and find strategies that lead to a potential treatment of COVID-19. Antibodies with highly diversified antigen binding specificity are produced from B-lymphocytes under V(D)J rearrangement that will define the antibody responses [5]. The variable structure in the heavy and light chains contain complementarity-determining regions (CDRs), where mutations are prone to occur. The CDR3 of the heavy chain covers the D gene and junction regions of D-J and V-D. The highly mutable CDR3 region could contribute to the diversity of antibodies, which is theoretically estimated at 10^15^ variants [6, 7]. Therefore, the sequence similarity analysis of CDR3s could infer the specificity of antibodies against antigens.

Four antibodies responsive to the earlier SARS-CoV-2 variant (WA-1) have been reported to be able to interact with a wide range of variants of concern (VOCs) [8]. The antibodies against different VOCs are readily found to suggest that antibodies with broad neutralization may exist [9–13]. On the other hand, convergent antibodies reactive to SARS-CoV-2 from different infected subjects suggest the diversified immune response against the SARS-CoV-2 may share similar clonotypes [3, 14]. It is useful to investigate naturally generated convergent antibodies from different time periods of the COVID-19 pandemic as infections of different variants constantly are appearing worldwide.

Bulk RNA-seq is a widely used methodology for the differential expression (DE) of genes [15–17]. Using routine DE analysis of RNA-seq can easily result in the loss of valuable information in the dataset. RNA-seq of lymphocytes using high throughput next generation sequencing provides a resource for mRNA transcript of antibody genes. Although single cell/bulk cell BCR-seq sequencing is available to retrieve the V(D)J sequences [7, 18], the abundant datasets of bulk RNA-seq are readily available while BCR-seq data are not generated. The immune repertoire reconstruction tools (for example: TRUST4, MixCR et al.) can help retrieve V(D)J sequences from Bulk RNA-seq [19, 20]. We have leveraged the TRUST4 to reconstruct the heavy chain VDJ for a longitudinal analysis of convergent antibodies.

## 2. Materials and Methods

### RNA-seq datasets

Four public RNA-seq data sets were retrieved from the Gene Expression Omnibus (GEO) [21]. Collectively, in these four data sets, there were 269 COVID-19 positive patients and 26 negative patients. These patients were distributed among 5 different groups (Figure 5a).

Samples in Group 1 were collected between March 2020 and April 2020. The data set consisted of 69 blood samples which were extracted from patients who were admitted to either the infectious disease unit or the designated ICU at a university hospital network in northeast France. All the patients in this group tested positive for SARS-CoV-2 through a qRT-PCR test which detected COVID-19 nucleic acids using a nasopharyngeal swab.

Samples in Group 2 were collected between April 2020 and May 2020 from patients that were admitted to either Albany Medical Center’s medical floor or the medical intensive care unit (MICU). The data set included 126 blood samples from 100 patients who tested positive for SARS-CoV-2 and 26 patients who tested negative for SARS-CoV-2.

Samples in Group 3 were collected between February 2021 and May 2021 by extracting blood from the buffy coat and was then purified with the Maxwell RSC simply RNA Blood Kit. In total, the data set included 100 blood samples which were extracted from 48 patients. We assigned 32 of the 48 patients to Group 3 as they had been classified as patients who had contracted the Alpha variant. Blood was extracted from patients through three samplings which took place 10 days, 25 days, and 45 days after the patient began feeling symptoms compatible to SARS-CoV-2. All 32 patients had their blood sampled 10 days after the patient started experiencing symptoms, 30 samples were collected 25 days after the patient started experiencing symptoms, and 5 samples were collected 45 days after the patient started experiencing symptoms. In our experiment, we combined each patient’s samplings into one sample.

Samples in Group 4 were from the same downloaded dataset as Group 3 but were classified as patients who had contracted the E484K escape mutation of the Alpha variant. There were 13 patients who tested positive for the mutation and samples were collected through three different samplings. All 13 patients were sampled 10 days after experiencing symptoms, 10 patients were sampled 25 days after experiencing symptoms, and 7 patients were sampled 45 days after experiencing symptoms. In this group, each patient’s samplings was combined as well.

Samples in Group 5 were collected between December 2021 and March 2022 from patients that were infected with the Omicron variant. Patients in this group were admitted due to either testing positive after attending an outpatient clinic at Krnakenhaus St. Vinzenz Zams using the SARS-CoV-2 PCR test or being in contact with a family member who had tested positive for SARS-CoV-2. 47 of the blood samples were collected zero, one, two, three, four, and five days after patients had tested positive for SARS-CoV-2 by the qRT-PCR test. 14 patients were sampled on day 0, 7 patients were sampled on day 1, 14 patients were sampled on day 2, 7 patients were sampled on day 3, 2 patients were sampled on day 4, and 3 patients were sampled on day 5.

### Reconstruction of VDJs and Longitudinal Analysis

The workflow is shown in Figure 1b. The RNA-seq raw reads from the same patient were merged into one fastq file and used as input for the bioinformatics tool TRUST4 [19, 22]. TRUST4 reconstructs immune repertoires and annotates the V(D)J assembly of each sample. Then, using the reconstructed immunoglobulin molecules, the variable regions, including the frameworks and hypervariable regions, are annotated.

**Figure 1.**
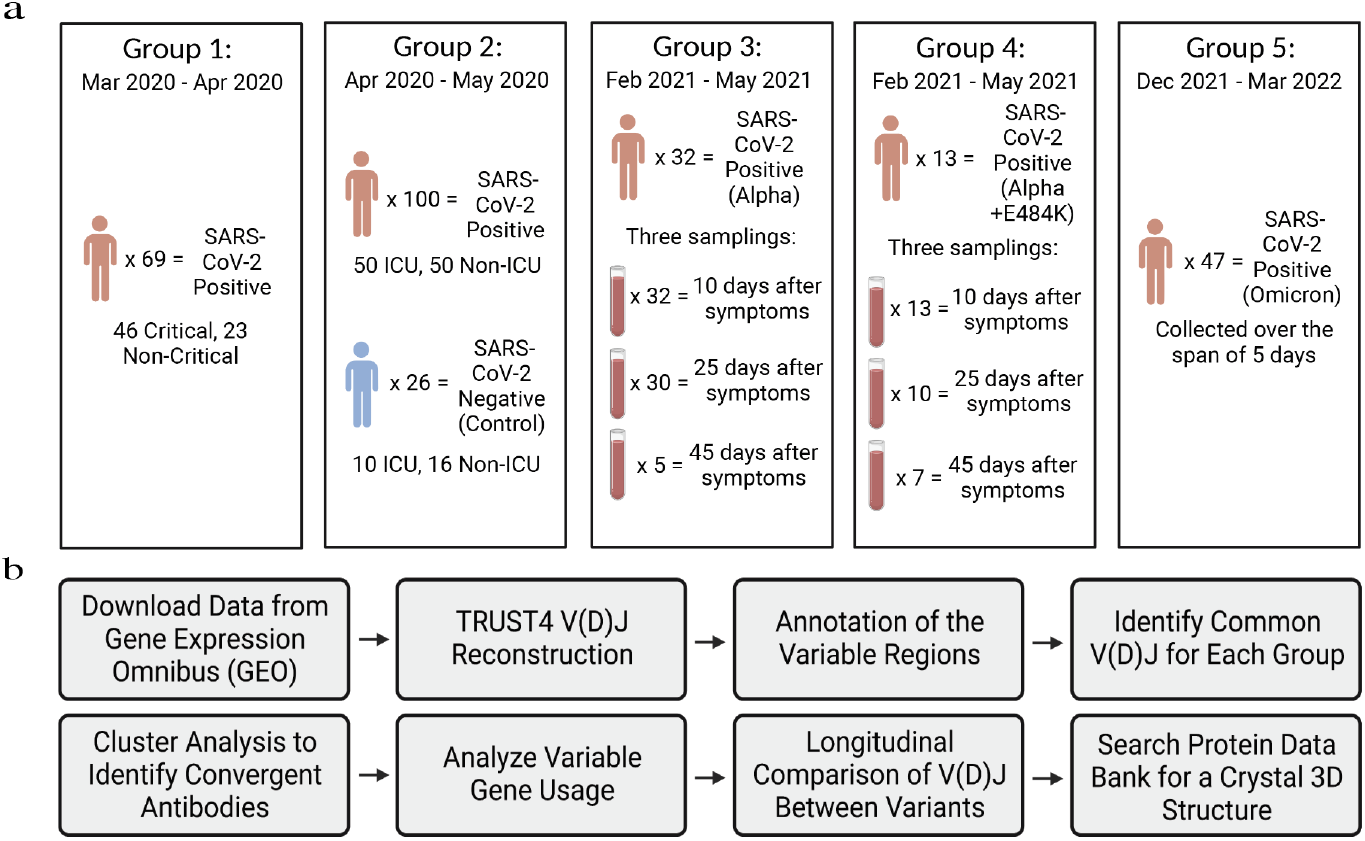
Overview of Sample Population and Experimental Design. (a) RNA-seq data from the same patient were combined in each group. All data were downloaded from the GEO databases (Group 1: GSE157103; Group 2: GSE172114; Group 3 and 4: GSE190680; and Group 5: GSE201530). SARS-CoV-2 variants in Group 1 and 2 were not identified. (b) The workflow of the longitudinal analysis of reconstructed V(D)Js.

The normalized score to evaluate the frequencies of common VDJs were calculated. Each group consisted of a different number of patients and a different number of VDJs predicted in each patient; therefore, the normalized score included the weight of the number of patients in each group. Thus, we could perform longitudinal comparison without the effects of each group’s population size. To calculate the normalized score of common VDJs we implemented the following equation:

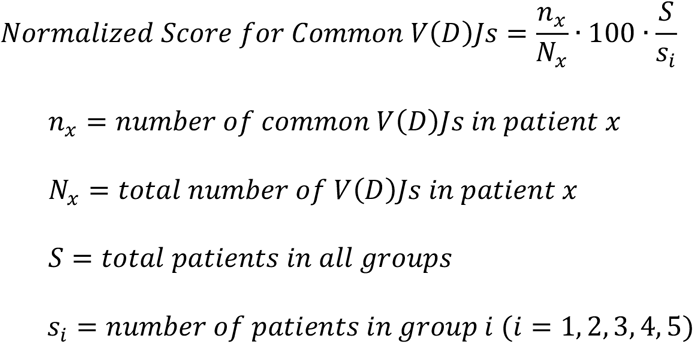

Nomalize scores for each patient are shown as violin plots in Figure 2b. The heavy chain VH gene usage was evaluated by the abundancy of each VH gene in the common VDJs. A heatmap was created to display the frequency of each VH gene in Figure 2c.

**Figure 2.**
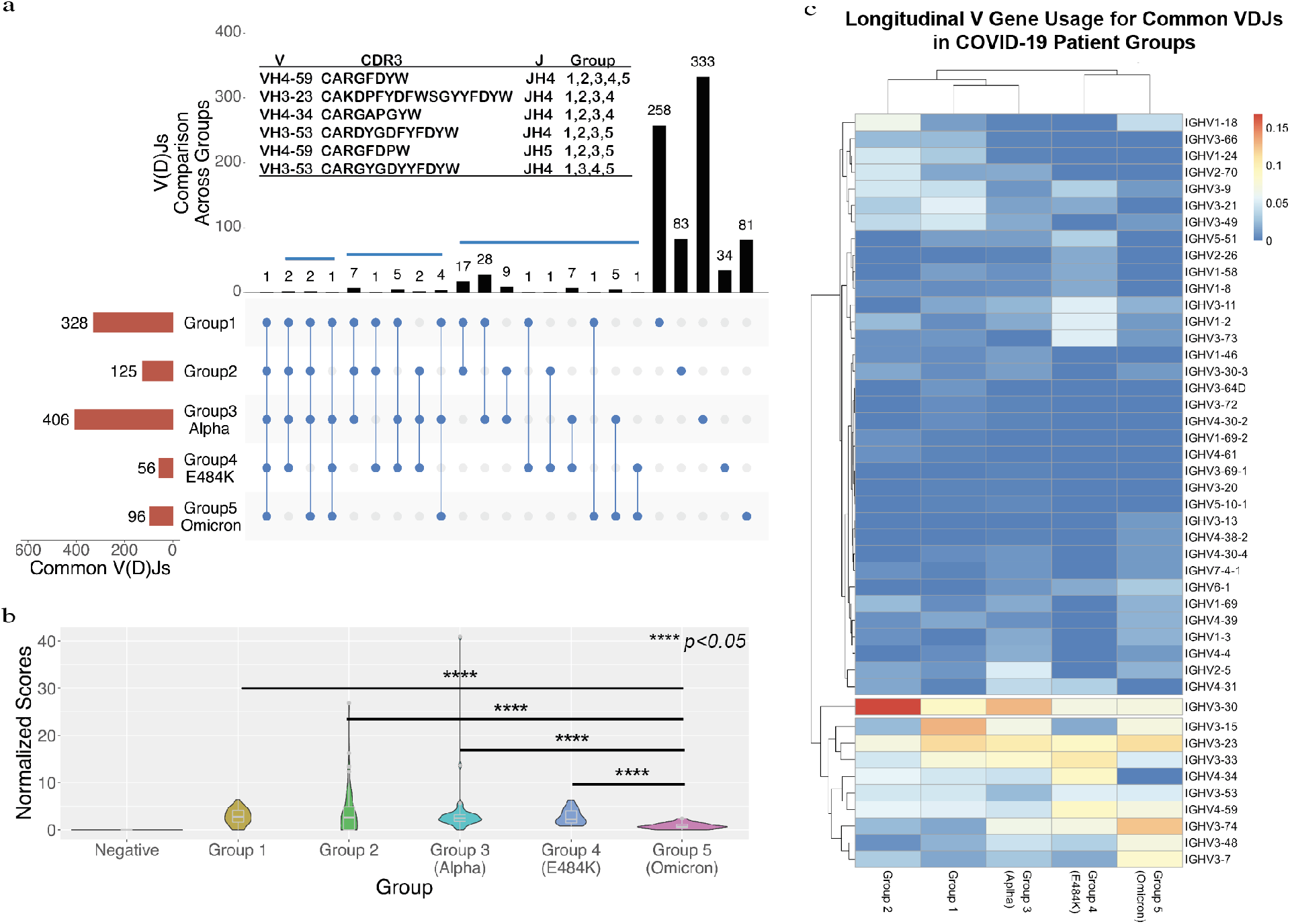
A Longitudinal Comparison of Common VDJs across the Five Different Groups. **(a)** The red bars at bottom left side show the total number of common VDJs for each group. The black bars at top show the number of IGC-VDJs and the number of common VDJs in only each group itself. The panel (a) also lists the V, CDR3 and J for the IGC-VDJs that were found in 4 or 5 groups. **(b)** Shows a comparison of common VDJs with the normalized scores. **** represents the p-value < 0.05 threshold using the one-tail t-test with heteroscedastic setting. **(c)** The heatmap of VH usage of VDJs between different groups. The frequencies of VH genes in common VDJs were utilized to generate the heatmap.

### Identification of convergent VDJ clusters and Protein Data Bank search with the reconstructed VDJ sequences

We performed a pair-wise calculation using the Levenshtein Distance [23] between CDR3 amino acid sequences from groups with the same VH and JH genes and the same amino acid sequence length of CDR3s. We set the Levenshtein Distance cutoff value to equal 2. The network was then visualized with the Gephi Software downloaded from (https://gephi.org). The common VDJ sequences were then used as queries to search the Protein Data Bank (PDB) to find the homologous VDJs from an antibody that was clinically proven to be against SARS-CoV-2.

## 3. Results

### 3.1. Identification of common V(D)Js from RNA-seq and the logitudinal analysis

There were 269 SARS-CoV-2 positive patients and 26 negative patients who served as a control group. Samples were grouped based on their SARS-CoV-2 variant type and/or the time they were collected. We reconstructed the V(D)Js using an immune repertoire reconstruction tool TRUST4 [19]. A total 629,137 heavy chain VDJs were reconstructed. The V, J and CDR3 regions were annotated based on the VDJ genes from the international ImMunoGeneTics information system (IMGT) [24, 25] and the human genome reference hg38. The immunoglobulins with common VDJs are defined as having the same V and J genes are used and the exact same CDR3 sequences are matched within each group. We identified a total of 1011 common VDJs in groups 1–5 as shown in Figure 2a. The common VDJs can be found in a diversified number of patients in each group (supplementary Table S1). The common VDJs were then analyzed in the longitudinal analysis. A common VDJ (VH4-59, JH4 and CDR3 sequence CARGFDYW) was observed in all five sample groups (Figure 1a). Five other VDJs were found across four out of the five groups. When comparing Group 3 (infected by Alpha variant) to Group 5 (infected by Omicron variant), we found 13 common VDJs that existed in both groups making up 2.7% of unique common VDJs in the two groups.

In order to compare the common VDJs longitudinally (Inter-Group Common VDJs as IGC-VDJs), the frequencies of IGC-VDJs in common VDJs of each group were adjusted by the number of patients in each group (see the calculation of normalized score in methods). The result shows a significant decrease in the normalized score for the patient group infected with the Omicron variant (Figure 2b), which may suggest that more patients tend to contain immunoglobulins with Omicron specific VDJs. The normalized score for the SARS-CoV-2 negative groups was zero further suggesting that the immunoglobulins with common VDJs found in other groups correlated with the viral infection.

To characterize the common VDJs further, we generated a heatmap of VH gene usages in all five groups. The VH3 gene family has been dominantly used in all groups. However, we observed that IGHV3-30 is more favorited in the variants from the early period of the COVID-19 pandemic such as Group 2 and Group 3 infected by the Alpha variant. IGHV3-74 and IGHV3-7 were preferred by Omicron-infected Group 5.

### 3.2. Identification and Verification of Convergent VDJs

The antigenic specificity of antibodies is mostly determined by their VDJ usage and diversified CDR3 regions [26]. Immunoglobulins with same V gene and J gene and similar CDR3 sequences have been suggested to respond to the same antigens [3, 14, 27]. Therefore, we are combining the CDR3 length and Levenshtein distance of amino acid sequences to characterize the CDR3 regions. If VDJs are using same V genes, J genes, same CDR3 length and Levenshtein distance is 2 or less, the VDJs will be classified as a convergent cluster.

We identified a total of 129 convergent clusters. Figure 3a visualized all the common VDJs and convergent clusters. The top 15 clusters with the most nodes are labeled. Clusters 2 and 15 covered all five SARS-CoV-2 positive groups (Table 1). We observed that Cluster 1, 2, 8, and 15 cover the Omicron-infected group and Alpha-infected groups. Cluster 3-7 and 9-14 were from groups that were not Omicron-infected (Table 1). The complete convergent VDJ list are available from the supplementary Table S2. The result indicates the possible variant specific antibodies existing in the groups.

**Figure 3.**
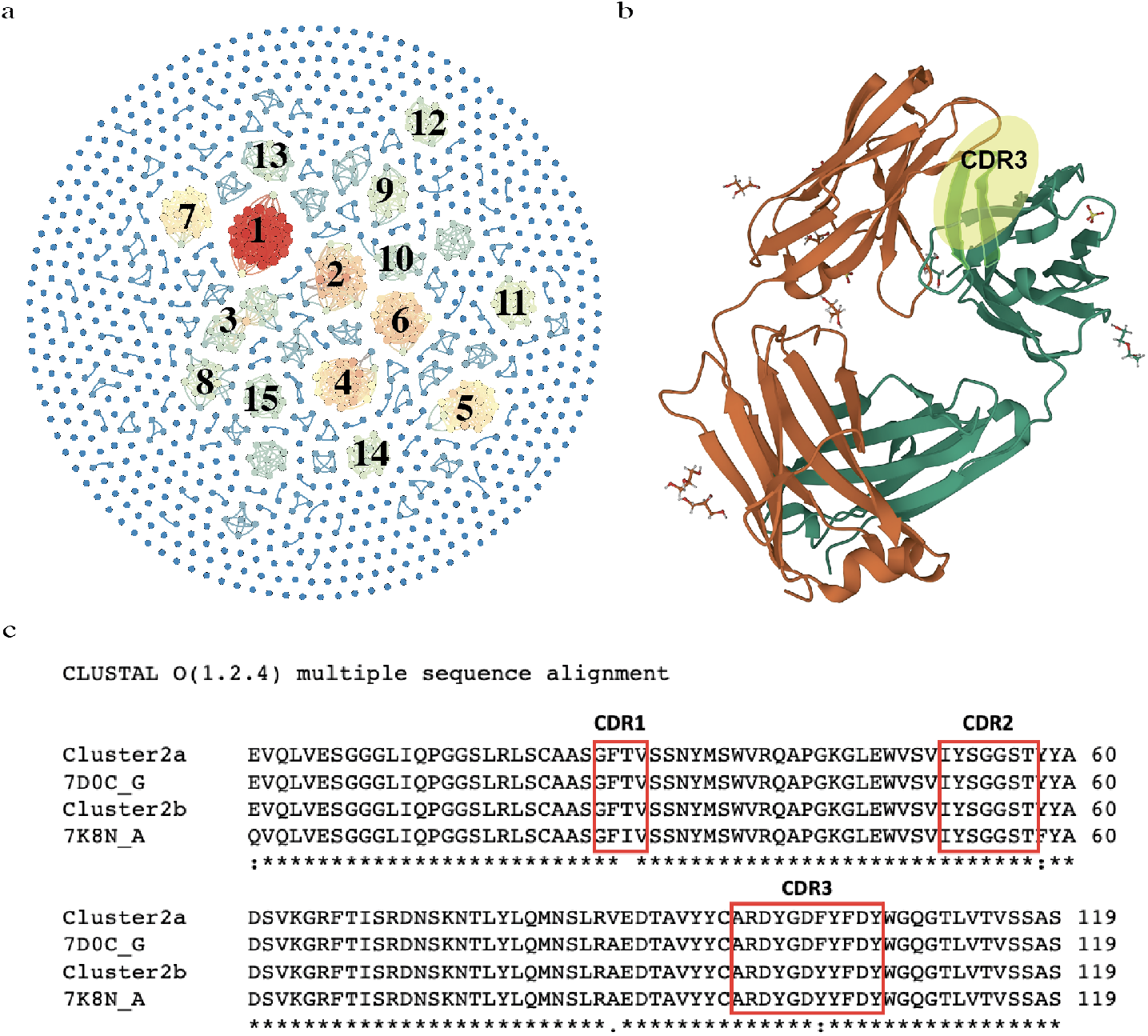
The identification of convergent VDJ clusters and comparison of the variable regions between the convergent cluster and the antibodies responsive to SARS-CoV-2. (a) The convergent clusters were colored based on the number of nodes in each cluster. (b) A Crystal Structure of an Anti-SARS-CoV-2 Human Neutralizing Antibody Fab Fragment (7K8N_A). The heavy chain CDR3 region is highlighted in yellow. (c) VDJ amino acid sequence comparison among two members of the convergent cluster 5, 7K8N_A and 7D0C_G from an antibody against SARS-CoV-2 in the Protein Data Bank (PDB).

To evaluate the convergent clusters, we leverage the Protein Data Bank (PDB) to search for a protein structure. The information of antibodies in PDB database may provide the antigenic specificity. We found that two Fab structures that are known to respond to SARS-CoV-2 have highly similar or the exact same CDR3 amino acid sequences to members of our identified convergent cluster 2 (Figure 3b and 3c) [28, 29]. The Fab fragment 7K8N is from a neutralizing antibody against the ACE2 receptor-binding domain (RBD) of SARS-CoV-2 [28]. 7D0C is from another neutralizing antibody that shows bivalent binding and inhibition of SARS-CoV-2 [29].

**Table 1.**
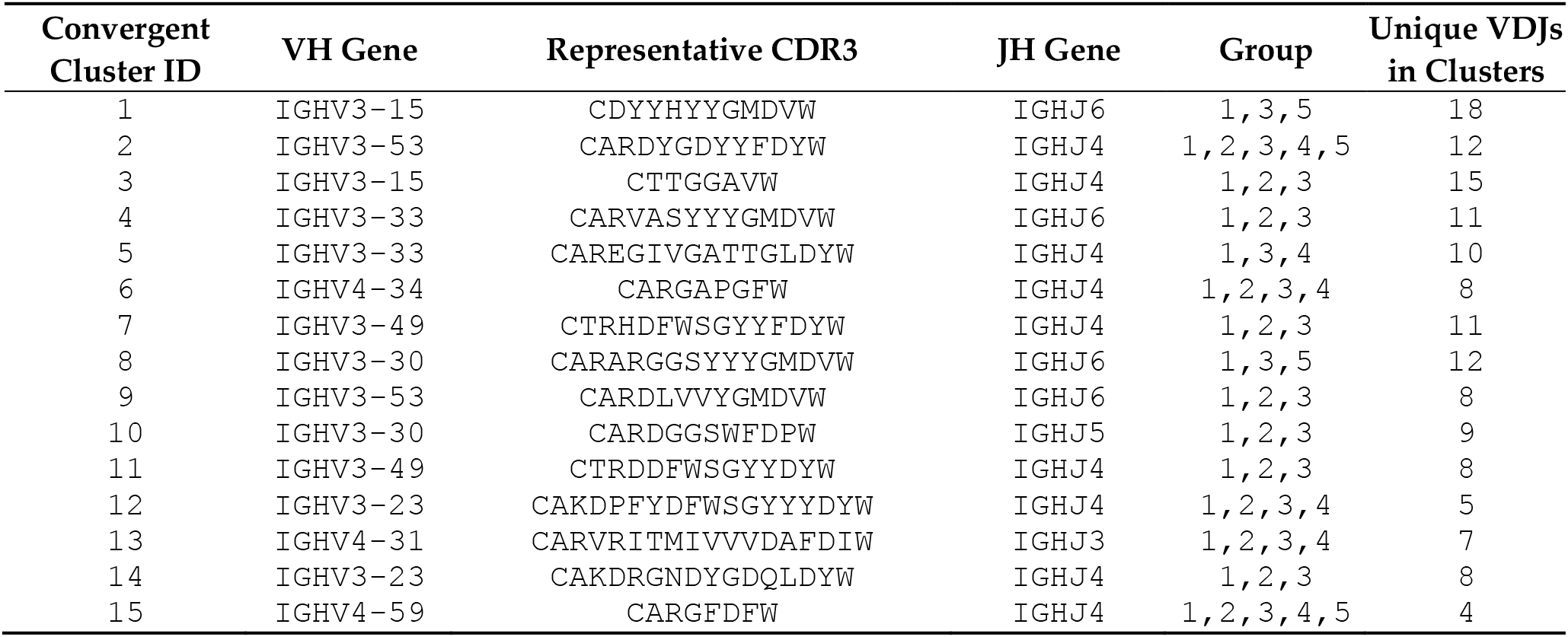
Representative CDR3 sequences, V genes and J genes for the top 15 Convergent VDJ clusters

## 4. Discussion

The convergent antibodies identified in this study have potential antigen specificity against SARS-CoV-2. The absence of common V(D)Js in the negative control group’s data set suggests the low antibody titer in normal blood serum and the very diversified VDJs are used for immunoglobulins generated by PBMCs. Our longitudinal study has shown that there are 2.7% common VDJs that exist in patients upon being infected with the Alpha or Omicron variants of COVID-19. The convergent analysis has suggested that the antibodies developed in different patients with similar antigenic specificity may potentially share similar clonotypes. Most of the convergent VDJ clusters are from multiple groups, which indicates that different SARS-CoV-2 VOCs occurring in different pandemic periods may induce similar clonality of B-lymphocytes [30]. This phenomenon has also been observed in the influenza virus vaccination [31]. The CDR3s of Cluster 7 and 11 show high similarity with one amino acid difference in the length of the sequence, which suggests that multiple clusters share similar antigenic specificity. In addition to the convergent CDJ clusters, our analysis result regarding the common VDJs show that most of the antibodies with common VDJs are group specific. The unique antibody profiling in each group suggests diversified immune response profiles were generated upon infection of SARS-CoV-2.

This project provides an efficient way to characterize the antibodies against SARS-CoV-2 in COVID-19 positive patients. The 3D structure of immunoglobulin Fab fragments in PDB database confirms that one convergent VDJ cluster in our study and provides validation for our analysis. Although the methods in the project were used to characterize antibodies against COVID-19, the procedure can easily be implemented to analyze other antibody-related immune responses against different viral and auto-immune diseases. This method allows us to identify the variable regions of the light chain as well. It is challenging to pair the heavy and light chain using bulk RNA-seq. However, a single cell RNA-seq can be processed for a just few samples and solve this challenge because the bulk RNA-seq aids with the process of identifying the individual samples containing the interesting antibodies. The specificity of convergent antibodies can be screened through the recombinant antibody production system.

## Supplementary Materials

Supplementary Table S1 and Table S2.

## Author Contributions

Conceptualization, K.J.L., M.A.Z and K.M.M.; Data analysis: K.J.L.; Writing—review and editing, K.J.L., M.A.Z and K.M.M.; supervision, K.M.M.; All authors have read and agreed to the published version of the manuscript.

## Data Availability Statement

The GEO accession numbers used in this study are Group 1: GSE157103; Group 2: GSE172114; Group 3 and 4: GSE190680; and Group 5: GSE201530

## Acknowledgments

The authors thank the technical advice from the bioinformatics core in the Department of Epigenetics and Molecular Carcinogenesis, The University of Texas MD Anderson Cancer Center; The authors acknowledge the support of the High-Performance Computing for research facility at the University of Texas MD Anderson Cancer Center for providing computational resources that have contributed to the research results reported in this paper. The authors acknowledge advice from the Recombinant Antibody Production Core (CPRIT RP190507). The authors also thank the research labs that deposited the RNA-seq data to GEO database and all the subjects enrolled in the original studies. The publication was supported by funds from UTMDACC.

